# Exercise on cytokine responses in males and females: Effect of leucine, HMB and BCAA

**DOI:** 10.1101/2023.07.11.548561

**Authors:** Alexander D. Brown, Maria Grammenou, Chris C.L. Tee, Mee Chee Chong, Claire E. Stewart

**Author notes:** Corresponding Author: Claire E. Stewart: Research Institute for Sport and Exercise Sciences, Liverpool John Moores University, Liverpool, UK ORCID: 0000-0002-8104-4819.

## Abstract

**Purpose:** To identify the effects of leucine, β-hydroxy β-methylbutyrate (HMB) and branched chain amino acid (BCAA) on post-exercise cytokine responses in females and males.

**Methods:** Males (n=53) and females (n=37) completed 100 drop jumps and consumed either no supplement, leucine (3g/d), HMB (3g/d) or BCAA (4.5g/d) from 1d pre to 14d post-exercise. Muscle soreness, squat jumps, chair rises and creatine kinase (CK) were measured at pre, post, 24h, 48h, 7 and 14d. Blood lactate (pre, post), 10 cytokines (pre, 24h, 48h, 7d) and oestradiol (pre, 7d) were also measured.

**Results:** Without supplementation post-exercise, soreness was induced in both males (6-fold) and females (5-fold). With supplementation, there were no increases in CK or oestradiol in females and no impact on muscle soreness, performance, or function in both sexes. In males, CK was elevated in untreated (48%) and leucine (69%) conditions vs baseline, but these were suppressed with HMB and BCAA. IL-7 was elevated in females vs males at baseline (6.3-fold), leucine increased IL-7 concentrations in females at 24h (17.0-fold), 48h (5.1-fold) vs males. With HMB, TNFr1-α increased in females at 24h (2.2-fold), 48h (2.3-fold) and 7d (2.3-fold) vs males. In males with BCAA, TNFr1-α decreased (P=0.06) from pre to 24h (6.8-fold), then increased (P<0.05) from 24 to 48h (8.0-fold).

**Conclusion:** Although supplements were without effect on soreness following exercise, the cytokine response was evoked by exercise and impacted significantly by leucine, HMB and BCAA in females vs males. This improved cytokine response in females could lead to improved resistance to damage.

## Introduction

Exercising improves health and fitness across the general population, especially in sedentary, obese and diabetic individuals (Kraemer and Ratamess 2005; O’Gorman and Krook 2011). Many exercise regimes involve eccentric contractions (Neufer et al. 2015) and can result in the delayed onset of muscle soreness (DOMS) and muscle inflammation (Hyldahl and Hubal 2014). These systemic changes promote recovery and adaptation and involve the secretion and homing of cytokines, including interleukin-1 beta (IL-1β), interleukin-6 (IL-6) and tumor necrosis factor - alpha (TNF-α) (Hyldahl et al. 2014). The supplements leucine, branched chain amino acids (BCAA) and β-Hydroxy β-methylbutyric (HMB) are commonly consumed in the sporting arena and literature reports benefits on stimulating muscle recovery and attenuation of DOMS (Howatson and Van Someren 2008; Owens et al. 2019). However, little is known of the effect of these supplements on the systemic environment following an acute bout eccentric exercise.

Cytokine responses are dependent on the intensity and duration of exercise, for instance, IL-6 and interleukin-8 (IL-8) were elevated following 6h running or marathon (Drenth et al. 1995; Suzuki et al. 2000) but not 5k running (Drenth et al. 1998). Further, increases in interleukin-10 (IL-10) and IL-8 were elevated following high intensity interval training (Dorneles et al. 2016). No differences in interleukin-4, interleukin-7 and IL-6 levels were evident following eccentric or concentric contractions (Hyldahl et al. 2014). All studies were performed in male participants. In middle aged men and women, following ∼50 km walking for 4 consecutive days, IL-6, -8, - 10, -1β and TNF-α were elevated compared to baseline and TNF-α and IL-1β were greater in males versus females over the duration of the study (Terink et al. 2018).

The supplement leucine has long been proposed as an essential amino acid for recovery and adaptation following exercise (Koopman et al. 2005; Wilson et al. 2009). BCAA and HMB are related to leucine, in that they either contain leucine (BCAAs), or are a metabolite of leucine (HMB). All three supplements are reported to reduce indices of muscle damage (Wilson et al. 2009; Howatson et al. 2012; Kirby et al. 2012) and promote muscle adaptation (Panton et al. 2000; Karlsson et al. 2004; Koopman et al. 2005). Leucine (250 mg/kg/d) has been shown to reduce creatine kinase (CK), but not soreness following 100 drop jumps in 10 young males compared with placebo control (Kirby et al. 2012). Whereas, BCAAs (20 g/d) attenuated CK, soreness and force impairment following 100 drop jumps in 6 trained males versus placebo controls (Howatson et al. 2012). Similarly, in 12 untrained females, following 140 squats, both muscle soreness and reductions in force were suppressed following 100 mg/kg/d BCAA ingestion (Shimomura et al. 2010). In contrast, two studies reported no effect of BCAA (7.3g for 2 days or 100 mg/kg/d) on maximal voluntary contraction (MVC) or CK levels following 120 lower limb extensions (Jackman et al. 2010) or electrical stimulation (Fouré et al. 2016) in young males. HMB (3 g/d) had no impact on force gain, soreness reduction or CK activity in young males that performed 55 knee extensions on both legs (Wilson et al. 2009). By contrast, HMB (3 g/d) before and for 48h post a resistance exercise session, decreased CK activity and perceived soreness in young males (Wilson et al. 2013). However, to date no individual study has investigated the effect of an acute bout of eccentric exercise in the presence of these supplements in males and females, at the level of cytokine responses.

Research on the recovery following acute exercise in females is sparse, with some studies reporting an effect of high oestrogen being capable of improving membrane stability, or altering anti-inflammatory properties, similar to changes observed with vitamin E (Kendall and Eston 2002). Women with high (270 pg/ml) compared to low (112 pg/ml) oestrogen concentrations had decreased CK following 30 minutes of downhill running, despite an increase in DOMS in both groups (Carter et al. 2001). However, post-menopausal females on hormone replacement therapy (oestradiol for 1 year) which equated to 255 pg/ml at the beginning of study did not reveal any differences in muscle torque or CK concentrations compared to those without hormone therapy following downhill running (Dobridge and Hackney 2004). In agreement, Sewright et al. (2008) also reported no effect of oestrogen in a cohort of females (n=58), with high oestradiol levels, following elbow extensors compared to males. Finally, only one study has investigated the impact of a progressive resistance training session on the immune response between males and females and reported greater granulocyte and lymphocyte activity and reduced CK activity in females vs males (Stupka et al. 2000). Interestingly, all the females used oral contraceptives and their oestrogen concentrations (60 pg/ml) were greater compared to males (7 pg/ml), however this had no effect on histological damage, but may possibly have elicited an effect on the altered immune cell recruitment (Stupka et al. 2000).

There is a distinct lack of data on the effect of protein supplements on female recovery following an acute bout of exercise and the subsequent cytokine responses. Therefore, the objectives of this study were to assess an eccentric intervention in females compared to males and also to examine the impact of leucine and its associated supplements on muscle soreness and cytokine responses. The hypotheses to be challenged were 1. acute eccentric exercise will cause DOMS and a loss in muscle function in females and males, 2. HMB and BCAA will improve DOMS and muscle function in females and males, in a manner similar to leucine, 3. HMB and BCAA will stimulate cytokines involved in muscle recovery, in a manner similar to leucine.

## Methods

### Participants and Ethics Statement

Following ethical approval, 90 healthy and recreationally active males (n = 53) and females (n = 37) voluntarily participated in the study. The participant characteristics and experimental groups are reported in Table 1. The exclusion criteria stated participants must not: 1. undergo any regular lower body strength training, 2. complete strenuous exercise that would cause unaccustomed eccentric damage, and 3. partake in the study if they experienced DOMS within the last 6 months. Following clear explanation of the protocol and exclusion criteria, the participants completed a health screening questionnaire and signed a consent form. The study was approved by Liverpool John Moores Research Ethics Committee (16/SPS/007) and was registered as a clinical trial (NCT04679519). The study protocol also adhered to the Declaration of Helsinki.

**Table 1.** Participant characteristics.

| Group | Sex | Number | Age (years) | Height (cm) | Weight (cm) |
| --- | --- | --- | --- | --- | --- |
| Control | M | 19 | 22 ± 2 | 178.0 ± 5.1 | 74.28 ± 8.33 |
|  | F | 10 | 23 ± 2 | 162.0 ± 6.2 | 58.25 ± 7.54 |
| Leucine | M | 16 | 22 ± 4 | 175.7 ± 5.4 | 71.59 ± 7.90 |
|  | F | 9 | 22 ± 2 | 163.3 ± 7.3 | 57.54 ± 9.57 |
| HMB | M | 9 | 23 ± 4 | 176.6 ± 7.1 | 75.40 ± 9.30 |
|  | F | 9 | 21 ± 2 | 164.3 ± 9.5 | 63.19 ± 13.07 |
| BCAA | M | 9 | 23 ± 4 | 175.4 ± 3.4 | 75.64 ± 10.39 |
|  | F | 9 | 22 ± 4 | 163.6 ± 7.3 | 63.86 ± 10.61 |

Participant's age, height and weight reported. No significant difference within sex in age, height and weight. Significant difference between sex in weight and height (all $P < 0.05$ ). Data displayed as mean ± SD.

### Experimental Design

Four independent groups (untreated, leucine, HMB and BCAA), included the minimum sample size (n=9: power=0.8: α = 0.05) which was determined based on previous literature revealing an increase in jump height at 24h (65.3 ± 5.2 cm vs 60.3 ± 3.3 cm) following BCAA ingestion (Howatson *et al*., 2012). Initial experiments were performed to assess feasibility and compliance to the supplementation. Therefore, initial studies focussed on 2 groups of males who were randomly assigned to untreated or leucine group. On ascertaining the intervention and supplementation were viable, the protocol was extended to include females and males who were randomly assigned to all groups (untreated, leucine, HMB and BCAA). The study period covered 15 days (Fig 1). The first day (day -1) was allocated to supplement preloading and familiarisation of all the measures (except blood collection). Baseline measures (pre), exercise protocol and post measures (post) were completed on day 0. For 14 days, the non-control participants ingested supplements, all completed muscle soreness measures and weighed food diaries. At the timepoints: pre, post, 24h, 48h, 7 days and 14 days post damage, the participants travelled to the laboratory and completed performance, functional measures and provided a blood sample (Fig 1). The measures included squat jumps, chair rises and soreness scores.

**Fig 1.**
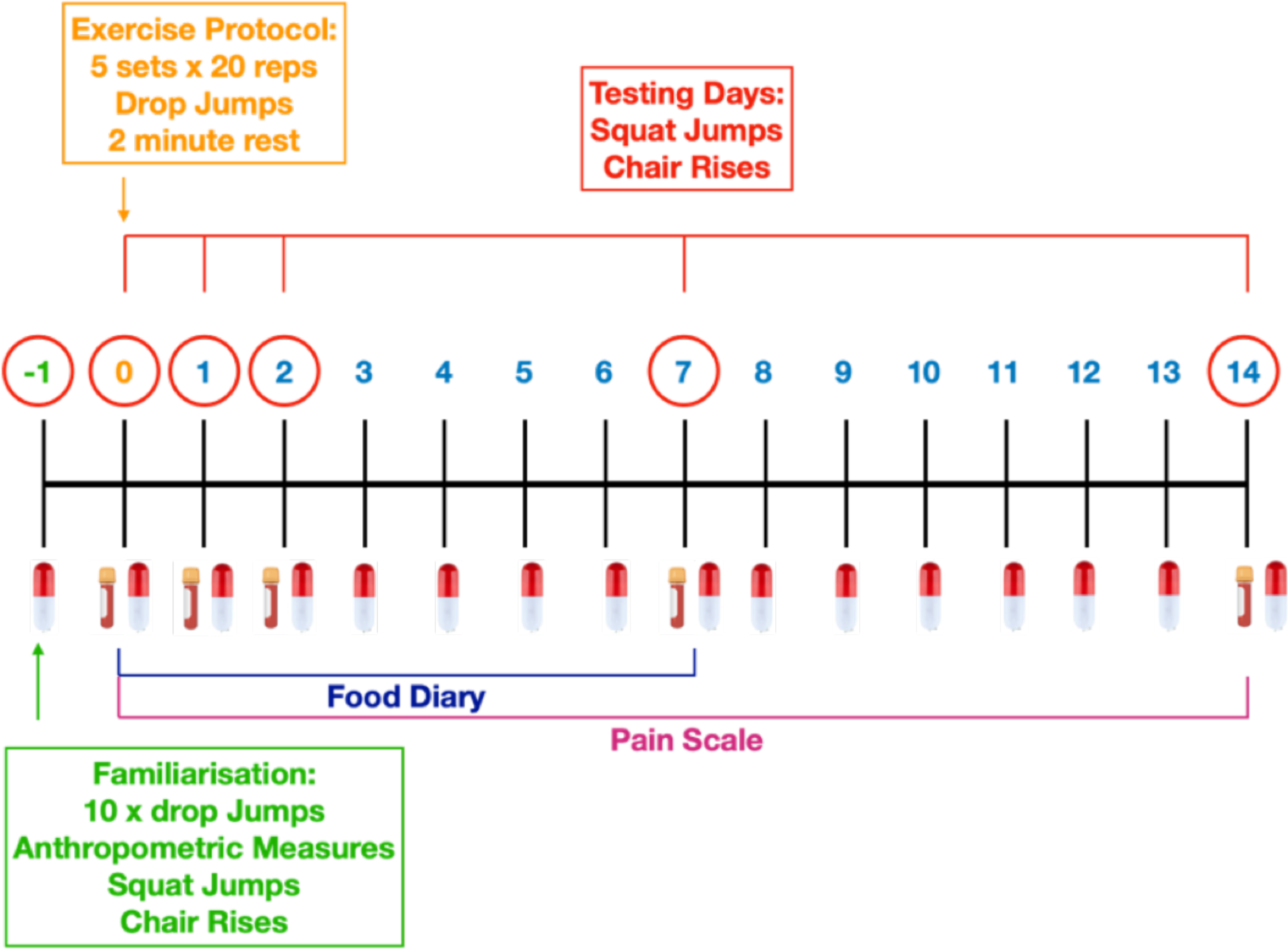
Schematic of the experimental design. Familiarisation occurred one day prior (D-1) to the exercise bout (D0). Supplements were ingested daily for 15D. On days (D0-1, 2, 7, 14) squat jumps and chair rises were completed and blood was collected. Food diaries from D0-D7 and soreness from D0-D14 were recorded.

### Anthropometric, Muscle Performance and Function Measures

During familiarisation, participants attended the laboratory and their body mass (kg), height (cm) and age (years) were recorded. Squat jumps were measured as an indicator of lower extremity power. The participants stood on a jump mat (Probotics, Inc, Huntsville, USA) with feet shoulder width apart and hands on hips. The participants lowered themselves into a squat position (knees approximately flexed to 90°) and held this position for 3 seconds, before being instructed to jump maximally, without arm movement. Squat jumps were performed three times and the average was recorded. Chair rises were performed to assess functional movement. The participants were asked to stand in front of a 40 cm box and stand with feet shoulder width apart and hands on hips. Participants were asked to sit on the box and stand back up, without arm movement. Participants executed as many chair rises possible within 30 seconds. The total number achieved was recorded. The participants were asked to rate their perceived muscle soreness of the lower extremity. They were provided with a visual analogue scale (VAS) with 0 = no soreness and 10 = worst possible soreness, and soreness was recorded daily.

### Supplementation Protocol

The participants either took no supplement or leucine, HMB or BCAA with doses selected based on safe, recommended daily allowance (RDA). For the leucine and HMB groups, this consisted of 3 g/d. BCAA tablets contained L-Leucine, L-Isoleucine, L-Valine at a ratio of 4:1:1, which consisted of 3 g/d leucine, 0.75 g/d isoleucine and 0.75 g/d valine. The supplements were purchased from Myprotein (Cheshire, UK) and taken in capsule form. The participants were asked to take either 1 (leucine and BCAA) or 2 (HMB) capsules with each meal (3 meals/day) to achieve appropriate dose. They were also asked to maintain habitual nutritional intake and to refrain from alcohol, ergogenic aids and anti-inflammatory medication. Habitual food intake was estimated via weighed food diaries over the first 7 days and data were analysed using Nutritics Software (Dublin, Ireland) with the participants’ calorie, carbohydrate, protein and fat reported (Table 2).

**Table 2.** Participant calorie and macronutrient intake.

| Group | Sex | Calories (Kcal/kg/d) | Carbohydrate (g/kg/d) | Protein (g/kg/d) | Fat (g/kg/d) |
| --- | --- | --- | --- | --- | --- |
| Control | M | 23.8 ± 3.2 | 2.7 ± 0.4 | 1.3 ± 0.2 | 0.9 ± 0.2 |
|  | F | 27.6 ± 4.4 | 3.4 ± 0.9 | 1.3 ± 0.4 | 0.9 ± 0.1 |
| Leucine | M | 26.5 ± 4.3 | 2.9 ± 0.5 | 1.3 ± 0.2 | 1.1 ± 0.3 |
|  | F | 27.0 ± 8.1 | 2.8 ± 0.9 | 1.4 ± 0.2 | 1.1 ± 0.5 |
| HMB | M | 26.1 ± 6.3 | 2.9 ± 0.7 | 1.4 ± 0.4 | 1.0 ± 0.3 |
|  | F | 24.3 ± 6.8 | 2.8 ± 0.7 | 1.3 ± 0.4 | 0.9 ± 0.3 |
| BCAA | M | 27.1 ± 7.9 | 2.9 ± 0.9 | 1.5 ± 0.5 | 1.1 ± 0.4 |
|  | F | 25.6 ± 5.2 | 2.7 ± 0.5 | 1.3 ± 0.2 | 1.0 ± 0.3 |

Participant's calorie, carbohydrate, protein and fat intake. Data normalised to individual body weight and daily amount averaged over the first 7 days. No significance between group or sex. Data displayed as mean ± SD.

### Eccentric Exercise Protocol

On day 0, the participants completed an eccentric exercise protocol. The protocol consisted of 100 drop jumps (5 sets of 20 reps), with a maximum of 10 seconds between reps and 2 minutes between sets. The participants were asked to stand on the 40 cm box, with hands on their hips and to drop off the box. The participants landed on a jump mat with feet shoulder width apart without moving hands. On impact, the participants immediately bent knee angle to approximately 90° and propelled themselves upward and landing with feet shoulder width apart. Each jump height was recorded, and participants were encouraged to maintain maximal effort and technical form.

### Blood Collection and Serum Separation

Blood samples were collected from the finger and the antecubital vein. At the timepoints pre, 48h and 7-day, samples were collected from the antecubital vein into 10 ml serum BD vacutainers and stored on ice for 30 minutes prior to centrifugation. Venepuncture was performed at these timepoints to allow for multiple analyses at timepoints corresponding to repair (48h) and recovery (7-day). At post, 24h and 14-day finger prick samples were collected into 500 µl microcuvettes and also stored on ice for 30 minutes prior to centrifugation and storage. For serum separation, all samples were centrifuged at 2000 rpm (800 g) for 10 minutes at 4°C.

### Blood and Serum Analysis

Blood lactate was measured pre and post exercise using a lactate strip and lactate analyser (Nova, Lactate Plus, Massachusetts, USA). CK was measured in all participants at all time points as follows: 20 µl serum was added to each well of a UV grade 96 well plate (San Jose, CA, USA), followed by 180 µl CK reaction reagent (Catachem Inc., Connecticut, NE). The plate was incubated for 5 minutes at 37°C. The change in absorbance was monitored continuously over 20 minutes using ELISA plate reader (Biotek, USA) at a wavelength of 340 nm.

Protein concentrations of multiple serum cytokines were measured using a CBA Kit (BD Biosciences: 558264) and flow cytometry, according to manufacturer instructions. The cytokines measured were: interleukins 1β, 2, 3, 4, 6, 7, 8, 10, 17α and tumor necrosis factor receptor 1 alpha (TNFr-1α). The timepoints measured were pre, 24h, 48h and 7 days. Briefly, premixed beads and PE conjugated antibodies were added to either standards or samples which were washed, resuspended and analysed using a BD Accuri C6 flow cytometer with BD CFlow^®^ Software, collecting 300 events per analyte (3000 events per sample). Following the manufacturers template, gating was applied to distinguish between each cytokine. Samples were analysed using Flow Cytometric Analysis Program (FCAP) Array software (BD Biosciences: v3). Calibration curves were plotted using 12 doubling dilutions from 2,500 – 0 pg/ml for all analytes except TNFr-1α, which was diluted from 10,000 pg/ml. 17beta-Estradiol concentrations were determined in female serum samples at pre and post 7 days using an ELISA kit (IBL International GMBH, Hamburg, Germany). Following the instructions provided by the manufacturers, standard, control and samples were pipetted into each well of a 96 well plate. The standards ranged from 0-2000 pg/ml and low (65-171 pg/ml) and high (244-642 pg/ml) controls were included. Subsequently, enzyme conjugate was added and the wells were washed, followed by the addition of substrate solution and finally, the stop solution before absorbance was quantified on an ELISA plate reader (Biotek, USA) at a wavelength of 450 nm. Concentrations of 17β-Estradiol in the serum samples were determined from the standard curve via linear regression calculation.

### Statistical Analysis

All data was tested for normality using the Shapiro-Wilk test. Dependent t-tests were utilised for comparisons between pre and baseline values. As expected, no statistical differences were identified and therefore, for time course comparisons, baseline data were not included. For all comparisons, three-way mixed ANOVA and Tukey post hoc pairwise comparison (to account for multiple variables tested simultaneously) tests were used. The between participant comparisons were between sex (male, female) and supplement (no supplement, leucine, BCAA, HMB). The within participant comparisons were over the time course (pre, post, 24h, 48h, 7d and 14d). Sample size and power analysis was performed using minitab (version 1.9) and statistical analyses were performed using GraphPad prism software (version 9). All data were represented as mean ± SD, unless mentioned otherwise.

## Results

### Participant Characteristics

There were no significant differences in participant height, mass or age within either female or male groups (Table 1). However, as expected, in each group females had significantly reduced (all P < 0.05) height and mass compared to males (Table 1). Nutritional data illustrated there were no significant differences in macronutrient intake in females and males (when normalised to mass) between any of the groups (Table 2). Protein intakes in females between the untreated (1.3 ± 0.4 g/kg/d), leucine (1.4 ± 0.2 g/kg/d), HMB (1.3 ± 0.4 g/kg/d) and BCAA (1.3 ± 0.2 g/kg/d) groups were not different. The same was true in males, with protein consumption in the untreated (1.3 ± 0.2 g/kg/d), leucine (1.3 ± 0.2 g/kg/d), HMB (1.4 ± 0.4 g/kg/d) and BCAA (1.5 ± 0.5 g/kg/d) groups, also not different. Notably, females and males consumed protein, on average, 0.5 g/kg/d and 0.6 g/kg/d greater than the RDA (0.8 g/kg/d) respectively.

### Examining the acute exercise protocol in the untreated group: females vs males

Blood lactate concentrations increased significantly from pre to post intervention in females (1.3 ± 0.6 - 3.6 ± 1.8 mmol/L: P < 0.05) and in males (1.2 ± 0.5 - 3.6 ± 1.8 mmol/L: P < 0.05) (Fig 2A), with no significant differences between sex, pre or post intervention (Fig 2A). There were significant increases in muscle soreness in females at post, 24h and 48h, all vs. pre intervention (4-fold, 5-fold, 5-fold respectively, all P < 0.05; Fig 2B). After which, levels of soreness decreased towards baseline from 48h to 7 and 14 days. As with females, there were significant increases in muscle soreness in males at post, 24h and 48h, all vs pre-intervention (4-fold, 5-fold, 6-fold respectively; all P < 0.05; Fig. 2B) and returned to baseline by 7 and 14 days. The increase in muscle soreness at 48h post exercise which decreased by 7 days was no different between females and males (Fig 2B).

**Fig 2.**
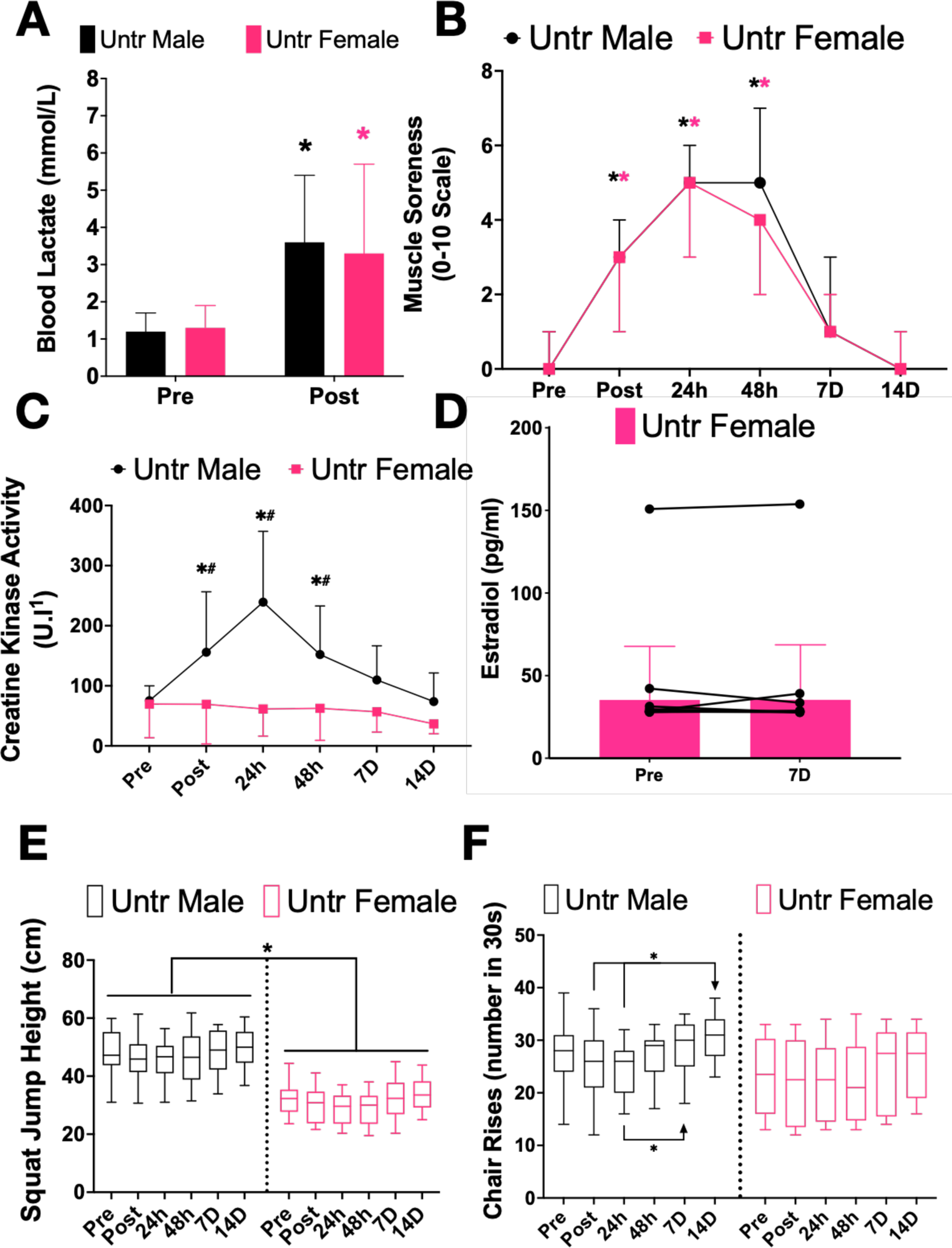
Impact of 100 drop jumps on blood and physiological markers between males and females. Blood lactate response pre and post exercise (A), significance in males (*) and females (*). Muscle soreness (B) from pre vs post, 24h and 48h, significance in males (*) and females (*). Creatine kinase activity (C) in males and females, significance between sex (*) and from pre to post, 24h and 48h (#). Oestradiol concentrations (D) in females at pre and 7-days post exercise. Squat jump height (E) over time, significance (*) between sex. Number of chair rises completed (F) and significance (*) represented from post, 24h to 14-days and 24h to 7-days. Data displayed as average ± SD. Significance set at P < 0.05.

Despite the increased muscle soreness in females, CK activity did not increase over time. In contrast to females, in males there were significant increases in CK in post, 24h and 48h time points vs. pre-intervention (2.1-fold, 3.2-fold, 2.1-fold respectively; all P < 0.05). Indeed, CK activity in females was significantly lower than in males at post (2.5-fold: P < 0.05), 24h (3.9-fold: P < 0.05) and 48h (2.4-fold) (Fig 2C).

Due to the anti-inflammatory properties and potential to improve membrane stability, high oestrogen concentrations are reported to attenuate CK response (Kendall and Eston 2002). However, all females bar one had circulating concentrations of <100 pg/ml, with oestradiol concentrations in the untreated at pre to 7-day being: 43.9 ± 40.4 to 44.0 ± 41.4 pg/ml (average ± SD) (Fig 2D).

Despite early increases in muscle soreness, there was no decline in squat jump performance following acute exercise in females or males (Fig 2E). Overall, males had on average, a significant increase (all P < 0.05) in jump height (47 ± 6.9 cm) compared to females (31 ± 4.8 cm) across time (Fig 2E).

The number of chair rises were quantified as a measure of muscle function. In females, increases in muscle soreness did not impact the number of chair rises performed across time (Fig 2F). In males, there were significant increases in the number of chair rises completed from post to 14D (1.2-fold: P < 0.05) and 24h to 7D and 14D (both 1.2-fold: P < 0.05: Fig 2F). Despite these differences, there was no significant difference in the number of chair rises completed between females and males across time (Fig 2F).

Together, these data suggest that 100 drop jumps in these participants increased muscle soreness but did not impact muscle function in females or males.

### Impact of supplements on muscle soreness and CK activity following exercise: females vs males

In an attempt to alleviate muscle soreness following 100 drop jumps, both females and males were supplemented with leucine, HMB or BCAA. Leucine was used as the internal control in these experiments to determine whether any differences were apparent between leucine alone or in combination with valine and isoleucine (BCAA with leucine concentration matched) or the metabolite of leucine, HMB.

Following 100 drop jumps, blood lactate significantly increased (all P < 0.05) in the leucine group from 1.2 ± 0.9 - 5.0 ± 2.6 mmol/L (4.2-fold), the HMB group from 1.1 ± 0.7 - 3.5 ± 1.5 mmol/L (3.2-fold) and the BCAA group from 1.3 ± 0.4 - 3.3 ± 1.3 mmol/L (2.5-fold) in females (Fig 3A). In males, in all the groups, from pre to post, there were significant (all P < 0.05) increases in blood lactate by from 1.4 ± 0.7 - 3.0 ± 1.6 mmol/L (2.1-fold) with leucine, from 1.5 ± 1.0-3.2 ± 1.4 mmol/L (2.1-fold) with HMB and from 1.5 ± 0.7-3.8 ± 2.1 mmol/L with BCAA (2.5-fold) (Fig 3A). There were no significant differences between supplementation groups or between sex at the post timepoint.

**Fig 3.**
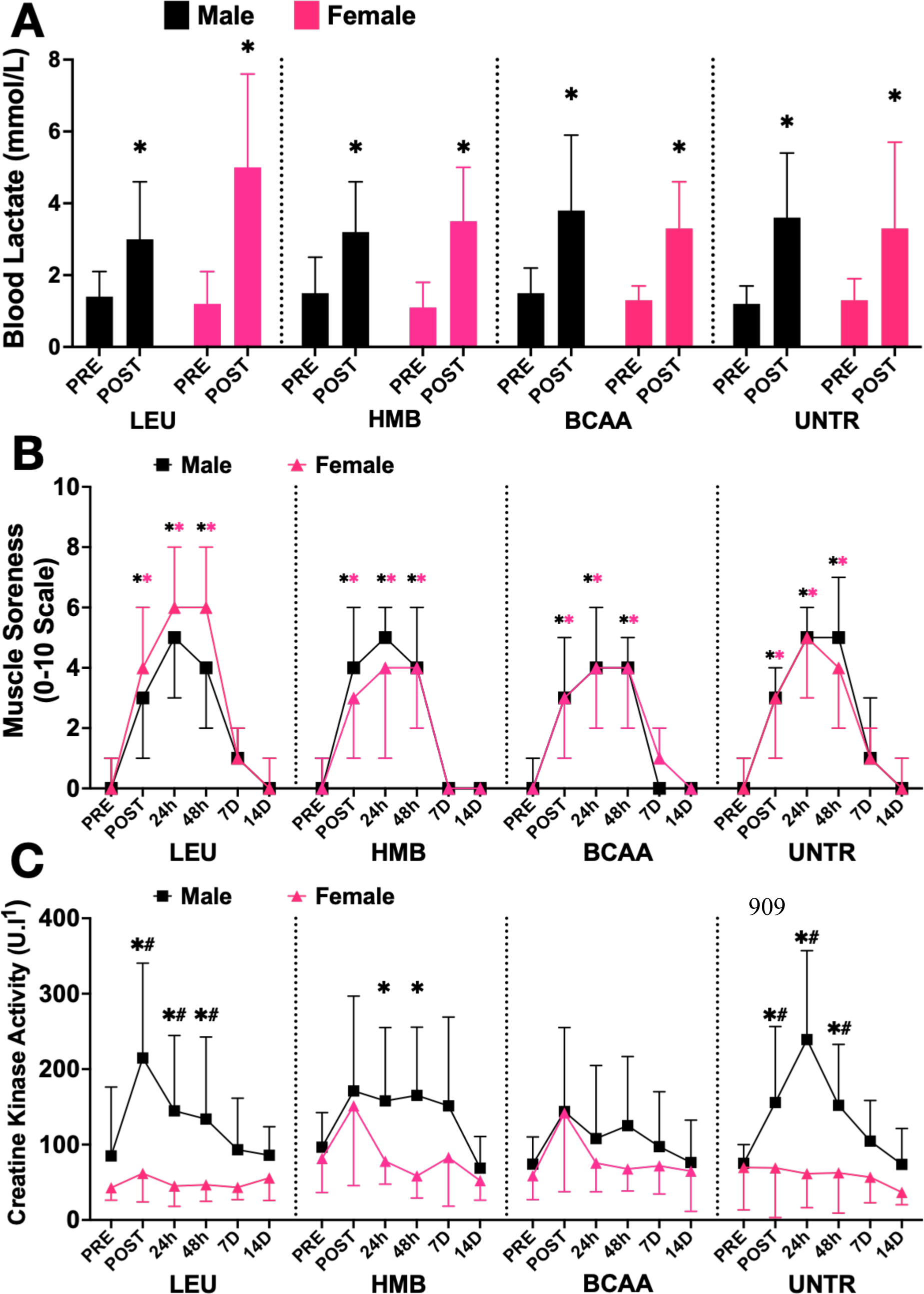
Effect of protein supplements on post exercise lactate, soreness and CK between males and females. Blood lactate response pre and post exercise (A), significance (*). Muscle soreness (B) from pre vs post, 24h and 48h, significance in males (*) and females (*). Creatine kinase activity (C) in males and females, significance between sex (*) and from pre to post, 24h and 48h (#). Data displayed as average ± SD. Significance set at P < 0.05.

There were significant increases in muscle soreness in all female groups, regardless of supplements (Fig 3B). In the female leucine group, there were significant increases (all P < 0.05) in pre vs post (4-fold), 24h (6-fold) and 48h (6-fold). In the female HMB group, significant increases (all P < 0.05) in pre vs post (4-fold), 24h (4-fold) and 48h (3-fold) were reported. Finally, in the female BCAA group, there were significant increases (all P < 0.05) in pre vs post (3-fold), 24h (4-fold) and 48h (5-fold). In all female groups, there were significant decreases in soreness scores from 48h to 7 and 14-day, with all values returning to baseline levels.

In all male participants, muscle soreness significantly increased from pre to post, 24h and 48h post intervention (all P < 0.05), regardless of the supplement group (Fig 3B). In the leucine group, there were significant increases in soreness (all P < 0.05) in pre vs post (5-fold), 24h (5-fold) and 48h (5-fold). In the HMB group, significant increases (all P < 0.05) in pre vs post (4-fold), 24h (5-fold) and 48h (4-fold) were reported. In the BCAA group, there were significant increases (all P < 0.05) in pre vs post (3-fold), 24h (4-fold) and 48h (4-fold). In all male supplement groups, soreness significantly decreased returning to baseline levels by 7 and 14 days (Fig 3B). There were no significant differences between females and males in muscle soreness responses, with trends following similar patterns (Fig 3B).

CK activity levels in the females did not increase over time in any group (Fig 3C). There were no differences between groups for CK activity values (Fig 3C). The female CK values per supplement group at onset (pre) of the intervention for leucine, HMB, BCAA and untreated groups were: 42.65 ± 16.37 U.l-1, 81.52 ± 45.12 U.l-1, 58.78 ± 31.71 U.l-1, 69.60 ± 56.10 U.l-1 respectively, and these are within reported normal ranges for female adults. Whereas, in males the basal CK concentrations in leucine, HMB, BCAA and untreated groups were: 84.94 ± 91.32 U. l-1, 96.51 ± 45.87 U. l-1, 74.13 ± 35.95 U. l-1, 75.24 ± 27.73 U.l-1 respectively, similar to those reported in females. In the male leucine group, there were significant increases (all P < 0.05) from pre to post (2.5-fold), 24h (1.7-fold) and 48h (1.6-fold). All CK values returned to baseline levels by 7 days in the leucine group. In contrast, the increase in CK activity in males supplemented with leucine, this increase was prevented in the presence of HMB and BCAA at all times, despite the reported increase in soreness (Fig 3B) and comparable increases in lactate reported across all male supplement groups (Fig 3A). Whereas there were significantly (all P < 0.05) reduced CK levels in females compared to males in the leucine group at post (3.5-fold), 24h (3.2-fold) and 48h (2.9-fold) and the HMB group at 24h (2-fold) and 48h (2.8-fold). There was no difference between females and males with BCAA supplementation.

Altogether, the supplements had no effect on muscle soreness in either females or males. However, HMB and BCAA, but not leucine, reduced CK activity in males and in the male BCAA group, CK was reduced to the activity levels in the female BCAA group. Given the potential impact of HMB and BCAA on reducing circulating CK levels, the impact of supplements on muscle function was investigated.

### Impact of supplements on muscle function and performance: females vs males

There was no impact on squat jump height with supplements or between groups (Fig 4A) in females. In addition, in males, there was also no difference with leucine, HMB or BCAA supplementation in squat jump performance over time (Fig 4A). As observed in untreated comparisons, in each supplement group, squat jump performance was greater in males compared to females. The numbers of chair rises completed by females were not significantly different across time in the leucine or BCAA groups (Fig 4B), whereas, in the HMB group, there were small albeit significant increases in chair rises completed from pre to 7-day (1.2-fold; P < 0.05) and pre to 14-day (1.3-fold: P < 0.05). Significant increases (both P < 0.05) were also observed from post to 7-days (1.2-fold) and post to 14-days (1.3-fold) in the HMB group only. Despite these within-supplement group changes, there were no significant differences compared to leucine or BCAA groups (Fig 4B) in females. In males, in the presence of leucine, chair rise numbers (Fig 4B) increased significantly (both P < 0.05) from post and 24h to 7-day (1.2-fold, 1.3-fold respectively). There were also significant increases in chair rise numbers from pre, post, 24h and 48h to 14-day (1.1-fold, 1.2-fold, 1.3-fold, 1.2-fold respectively; all P < 0.05). By contrast, in males, HMB and BCAA treatments were without significant impact on chair rises completed across the whole study period, despite the apparent reduction in CK activity. In addition, there was no difference between females and males in the number of chair rises performed with either supplement consumed (Fig 4B).

**Fig 4.**
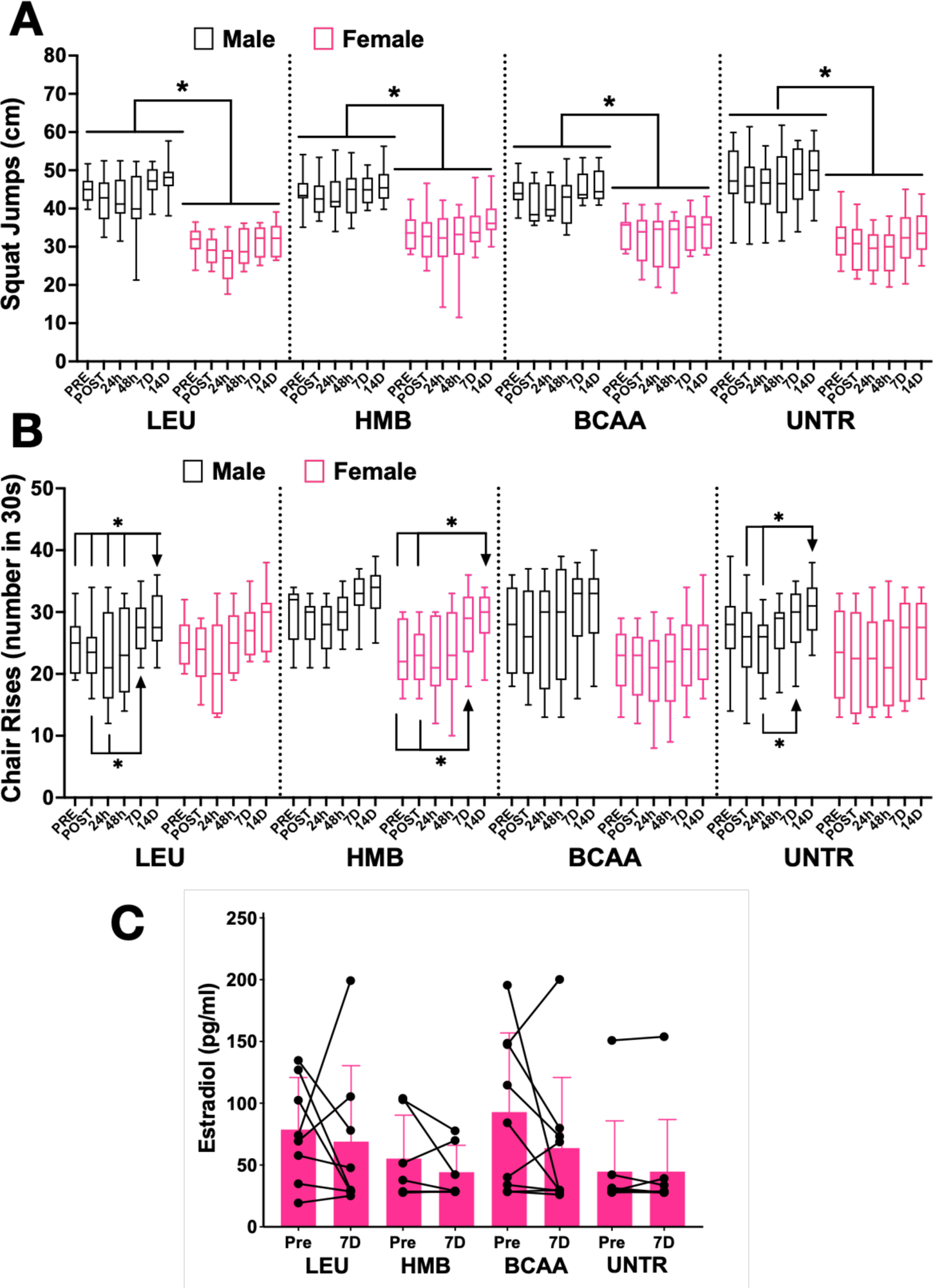
Effect of protein supplements on post exercise physiological measures between males and females. Squat jump height (A) over time, significance (*) between sex in the presence and absence of supplements. Number of chair rises completed (B) and significance (*) represented from pre, post, 24h, 48h to either 7 or 14-days. Oestradiol concentrations (C) in females at pre and 7-days post exercise. Data displayed as average ± SD. Significance set at P < 0.05.

The above differences between females and males and lack of impact of supplements following an acute bout of soreness inducing but non damaging exercise could be influenced by high oestrogen concentrations. However, there were equivalent concentrations of oestrogen in all female groups, therefore, the above measures were not impacted by changing oestradiol concentration (Fig 4C). The average ± SD oestradiol concentrations from pre to 7 days in the leucine, HMB and BCAA group were; 77.4 ± 41.4 to 67.8 ± 60.4 pg/ml; 54.3 ± 34.5 to 43.5 ± 21.4 pg/ml; 91.3 ± 63.0 to 62.7 ± 56.1 pg/ml, respectively. Individually, although there were fluctuations in oestrogen concentrations; that is a decrease or increase from pre to 7D, none of the concentrations were above 200 pg/ml. Further, there was no relationship between any individual participant who had higher concentrations at pre or 7D with CK or cytokine concentrations.

### Sex differences in cytokine concentrations with supplementation

From the 10 cytokines selected for measure, IL-3 and IL-10 were undetectable in all samples, IL-1β, 2, 4, 6 and 17 were all detectable but were not significantly different across any measures (Supplementary File). However, IL-7, IL-8 and TNFr-1α showed significant adaptations (Fig 5A-C). In the presence of leucine supplementation, there were significantly higher (all P < 0.05) concentrations from 0.57 ± 0.18 pg/ml in females compared to 0.09 ± 0.07 pg/ml in males (6.3-fold) pre-exercise intervention. This difference was enhanced at 24h from 0.68 ± 0.16 pg/ml in males vs 0.04 ± 0.04 pg/ml in females (17.0-fold) and returned to pre-value differences at 48h, 0.72 ± 0.22 pg/ml vs 0.14 ± 0.11 pg/ml (5.1-fold) (Fig 5A). These data were surprising given no impact of leucine on biochemical markers, function or soreness in females. There was no impact of HMB or BCAA supplementation on IL-7 concentrations in females or males across time (Fig. 5A). Untreated female IL-7 levels were unchanged without supplementation. In untreated males IL-7 concentration tended to increase with time, although significance was not achieved (Fig. 5A). Nevertheless, at 7 days, IL-7 concentration was significantly greater (P < 0.05) from 1.35 ± 0.43 pg/ml in males to 0.24 ± 0.12 pg/ml in females (2.6-fold; Fig 5A).

**Fig 5.**
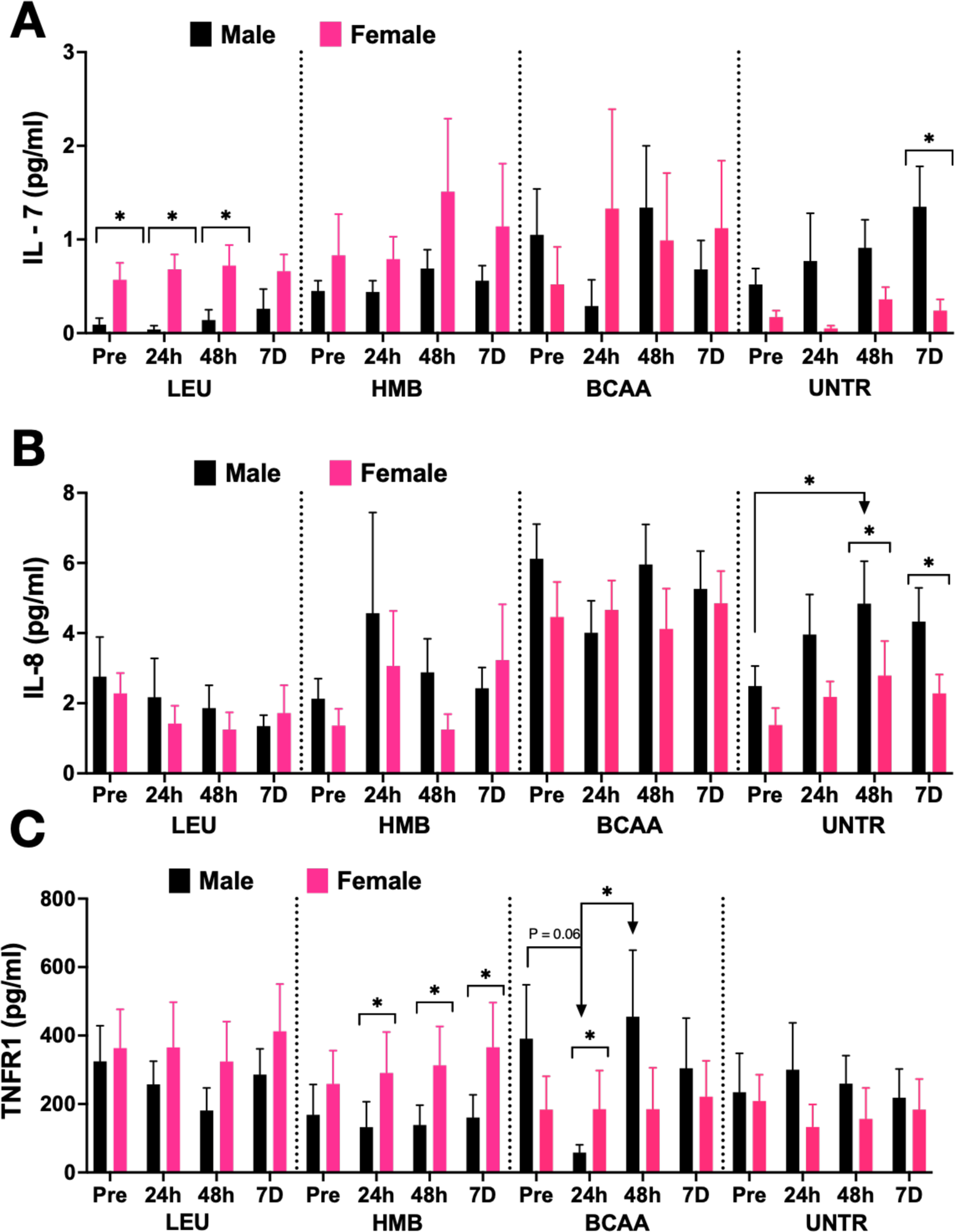
Effect of protein supplements on post exercise cytokines between males and females. Concentrations of IL-7 (A), IL-8 (B) and TNFr-1α (C) response pre and 24h, 48h and 7-days post exercise in the presence and absence of supplements, significance between sex (*) and between time indicated with an arrow. Data displayed as average ± SEM. Significance set at P < 0.05.

In contrast to IL-7 and in the presence of leucine, IL-8 concentrations remained at baseline levels for all time points and in both sexes (Fig 5B). In the presence of HMB, IL-8 levels were variable, with no differences evident over time or between sexes, again. Furthermore, BCAA supplementation tended to result in higher IL-8 concentrations in both males and females compared to other groups, however significance was not attained, and values were not significantly different from baseline (Fig 5B). Similar to IL-7, under control conditions, IL-8 concentrations increased over time in males (Fig 5B). Significance was reached from pre to 48h with concentrations from 2.49 ± 0.57 pg/ml to 4.84 ± 1.21 pg/ml (2.0-fold: P < 0.05). In addition, there were significant elevations (both P < 0.05) in males vs females at both 48h, with concentrations from 4.84 ± 1.21 pg/ml vs 2.79 ± 0.98 pg/ml (1.8-fold) and 7-day, 4.33 ± 0.96 pg/ml vs 2.28 ± 0.54 pg/ml (2.0-fold).

Finally, TNFr-1α concentrations were not different in the leucine group over time or between sexes, which was in contrast to IL-7 and IL-8 responses (Fig 5C). With HMB supplementation there were no differences in males over time. However, there were significant increases (all P < 0.05) in females at 24h; 290.71 ± 119.62 pg/ml vs 132.52 ± 74.02 pg/ml (2.2-fold), 48h; 312.71 ± 114.05 pg/ml vs 138.88 ± 57.68 pg/ml (2.3-fold), and 7 days; 365.79 ± 130.52 pg/ml vs 160.39 ± 66.28 pg/ml (2.3-fold) compared to males. In males with BCAA supplementation, TNFr-1α decreased (6.8-fold: P = 0.06) from baseline to 24h, and then significantly increased by 48h; 57.93 ± 22.71 pg/ml vs 455.17 ± 194.51 pg/ml (8.0-fold: P < 0.05) back to baseline (Fig 5C). There were no changes in females compared to baseline, however, at 24h, TNFr-1α concentrations were significantly elevated compared to males; 184.76 ± 113.16 pg/ml vs 57.93 ± 22.71 pg/ml (3.2-fold: P < 0.05). Similar to leucine, there was also no difference in TNFr-1α concentrations between sex across time.

In summary, with leucine and HMB supplementation there were increases in IL-7 and TNFr-1α concentrations in females vs males. BCAA supplementation inhibited the TNFr-1α response at 24h in males. Under un-supplemented conditions in males, IL-7 and IL-8 concentrations were elevated over 7 days compared to females. Interestingly, the changes in cytokines did not appear to associate with physiological, biochemical or soreness measures, however, may be important indicators of inflammation.

## Discussion

The objectives were to assess an eccentric intervention in females versus males and to examine the impact of leucine, HMB and BCAA on muscle soreness and cytokine responses. The protocol induced elevated lactate and muscle soreness (nociceptor activation in the extracellular matrix (Hyldahl and Hubal 2014)), in both sexes. In females only, this did not induce an increase in CK activity (measure of plasma membrane permeability), corresponding to previous studies (Stupka et al. 2000; Sewright et al. 2008; Shimomura et al. 2010; Hicks et al. 2016) and was not impacted by oestrogen. Surprisingly, there was no impact on muscle performance or function measures, despite the occurrence of DOMS in both females and males.

The cytokine panel investigated included many proteins that are involved in muscle inflammation, regeneration and adaptation (Paulsen et al. 2012). For instance, IL-6 is reported to mediate both pro- and anti-inflammatory responses following exercise and increases following downhill running, 6k run or marathon racing (Drenth et al. 1995; Suzuki et al. 2000; Nieman et al. 2001). Other pro-inflammatory cytokines included IL-1β, TNF-α, IL-17α and IL-8, which are reportedly increased following muscle damage and anti-inflammatory cytokines that mediate regeneration and adaptation; IL-4, IL-10 are also reportedly increased with exercise (Paulsen et al. 2012). Furthermore, Haugen et al. (2010) reported that IL-7 is produced by primary muscle myotubes and is reported to activate T-cells which are important during the inflammatory process. IL-7 levels were also increased in wounded keratinocytes and improved their migration (Bartlett et al. 2016). Finally, IL-2 and IL-3 modulate the immune system in response to infection and inflammation (Abbas et al. 2018; Dougan et al. 2019).

In response to the eccentric protocol or supplements there was no elevation in the majority cytokine concentrations coinciding with concentric versus eccentric contractions in males (Hyldahl et al. 2014). The induction of DOMS in the absence of severe damage may underpin this lack of elevation in these cytokines. In our study, the IL-7 profiles were altered with leucine supplementation, IL-7 concentrations increased in females versus males whereas with HMB and BCAA supplementation there was no difference between sexes. In males without supplementation, there was an exponential increase in IL-7, which reached significance at day 7 compared to females. One study also demonstrated increases in IL-7 immediately and 30 min post an acute whole body resistance training session in males (Kraemer et al. 2014). In contrast to these data, there was no change in IL-7 concentrations (Hyldahl et al. 2014), however, evidence from cell models does indicate that higher levels of IL-7 support immune cell recruitment, cell migration and potentially accelerated recovery, which could be important for females who supplement with leucine.

IL-8 is also reported to regulate leukocyte cell migration during wound healing (Engelhardt et al. 1998). Our results suggest that leucine increases IL-8 in females compared to males, with no impact of HMB or BCAA. One study investigating the impact of a resistance training session in a placebo versus HMB supplemented group of males also demonstrated increases in IL-8 at 30 min (Kraemer et al. 2014). In our study, the untreated male group exhibited increased concentrations of IL-8 versus females, which peaked at 48h, similar to the CK profile and following high intensity interval training in males (Dorneles et al. 2016). However, there was no difference in IL-8 concentrations between middle aged men and women following ∼50 km walking (Terink et al., 2018). Other work also revealed no change in IL-8 concentrations immediately, 1h, 3h, 6h, 24h, 48h, 72h and 96h post dumb bell exercises in men without supplementation (Hirose et al. 2004).

TNF-α is produced by macrophages and T-cells and has multiple synergistic roles in the removal of necrotic tissue, skeletal muscle satellite cell activation and proliferation (Warren et al. 2002). In addition, circulating levels of the soluble TNF-α receptor, TNFr-1α, correlate with increases in TNF-α post injury (Zádor et al. 2001; Liu and Tang 2014) and both are important in the inflammatory response during repair (Zádor et al. 2001; Fu et al. 2015). We reported no impact of leucine on the TNFr1-α response between sexes, whereas, with HMB supplementation increases were evident in females versus males. In this study, at 24h in males there was a significant repression of TNFr1-α compared to pre, which recovered to baseline by 48h. Townsend *et al*. (2013) also reported an attenuation of TNFr1-α with HMB (3 g/d) supplementation, following a resistance exercise training session in males. In contrast, with no supplementation and with HMB there was no difference in TNF-α concentration following a resistance training (Kraemer et al. 2014). TNFr1-α concentrations in the untreated group were not different between sex and did not change over time, which contrasts with Terink *et al*., (2018) where males exhibited greater TNF-α concentrations throughout.

High oestrogen concentrations can impact muscle performance, function and recovery (Kendall and Eston 2002). There was no difference between groups and individual responses indicated that most participants had low oestradiol concentrations (78% below 100 pg/ml) at both pre and 7d. Oosthuyse et al. (2017) reported in females at varying stages of their menstrual cycle versus males that there was no association between oestradiol concentrations and muscle soreness and CK. Alternatively, the differing phases of the menstrual cycle could affect the stiffness of the muscle-tendon unit in female participants (Eiling et al. 2007), thereby reducing or increasing susceptibility to muscle damaging protocols, dependent on the tendon stiffness. Indeed, some evidence indicates that CK concentrations following downhill running was significantly lower in females with higher (270 pg/ml) versus lower (112 pg/ml) oestrogen concentrations (Carter et al. 2001). Therefore, higher levels of oestrogen during menstrual cycle could have protective effects against muscle damage (Kubo et al. 2009). However, studies that report no impact of menstrual cycle on tendon mechanical properties (Burgess et al. 2009; Kubo et al. 2009) and on CK or soreness (Miles and Schneider 1993; Sorichter et al. 2001).

The majority of literature assessing the impact of nutritional interventions in eccentric exercise has compared protein supplements to a placebo/control following muscle damage in males (Wilson et al. 2009; Jackman et al. 2010; Howatson et al. 2012; Kirby et al. 2012), not females and frequently at exercise levels that evoke severe, debilitating damage. With this in mind, we utilised an intervention that would induce soreness, similar to more conventional (vs. laboratory-based) exercise, in the absence of severe damage in males and females enabling determination of the impact of leucine (experimental control) vs HMB (metabolite of leucine) or BCAA (leucine plus valine and isoleucine) supplementation following acute exercise in females and males. In both sexes, blood lactate levels significantly increased from pre to post with no differences evident between groups or sex. This suggests that all participants worked at comparable intensities throughout the exercise protocol.

In addition, muscle soreness significantly increased in both females and males with leucine supplementation over 48h, then reduced at 7 and 14d, with no beneficial impact of BCAA or HMB. Previous evidence comparing females and males with BCAA supplementation reported attenuation in female soreness compared to males following 7 sets of 20 squats (Shimomura et al. 2010), whereas, another study reported no impact of milk supplementation on 6 sets of 10 repetitions of hamstring contractions (Rankin et al. 2015). In males, attenuation of muscle soreness with HMB and BCAA supplementation compared to placebo has been reported (Jackman et al. 2010; Howatson et al. 2012; Wilson et al. 2013). Whereas no attenuation with leucine supplementation (Kirby et al. 2012) has also been reported. These contradicting findings are likely attributed to the dose of supplement, which includes 250 mg/kg/d (Kirby et al. 2012), 3 g/d (Wilson et al. 2009), 7 g/d for two consecutive days (Jackman et al. 2010) and 20 g/d (Howatson et al. 2012). Furthermore, the habitual protein intake should also be considered, and although subjective methods for measuring protein intake have limitations, protein intake above the current RDA of 0.8 g/kg/d, as in this study, likely contribute to differences in the literature.

CK concentrations did not change in females any group over time and in males, CK increased coinciding with increases in muscle soreness under control, leucine and HMB supplementation conditions. There was no difference immediately following exercise between sexes in the HMB group. In contrast, CK concentrations were attenuated in males to that of female levels with BCAA supplementation, despite DOMS occurring in both sexes. Shimomura et al. (2010) also reported no difference in CK with BCAA supplementation (100 mg/kg/d) in females. The increase in soreness but not CK could be due to the reduced load (body mass) the females encountered during the drop jumps compared to the males, which was perhaps insufficient to disrupt the cell membrane. Moreover, in support of our BCAA-supplemented CK data, Rankin et al. (2015) reported no difference in either males or females in CK activity over 72h with 6 sets of 10 repetitions of hamstring contractions with milk supplementation. The absence of a leucine response compared to BCAA is somewhat surprising and little evidence reports isoleucine and valine independently from BCAA which could offer potential unexplored insights.

Despite the DOMS, our results suggested no decline in muscle performance, following the exercise intervention protocol. However, Shimomura et al. (2010) suggested that BCAA supplementation attenuated maximal voluntary contraction (MVC) in females, however, our study did not determine MVC as an output marker of force production (Damas et al. 2016). Nevertheless, disparities in results could have been due to single joint knee extensions that were measured (Shimomura et al. 2010) were more sensitive to neural changes rather than functional changes. Also, jump performance utilises both legs simultaneously spreading the production of force compared to single leg measurements. Previous research also suggests that after eccentric contractions, untrained males and females display similar relative responses of muscle function (Hicks et al. 2016). Our results support this evidence, in that there were no differences between males and females in muscle function in the absence or presence of leucine, HMB or BCAA.

To conclude, comparison between sexes, using an intervention which induced soreness, indicated that females had attenuated CK levels compared to males, although soreness levels were not different. BCAA supplementation attenuated the increase in CK in males to that of females. Cytokine responses (IL-7, IL-8, TNFr1-α) were altered with leucine, HMB and BCAA supplementation, although in the untreated group, males had increased cytokine (IL-7, IL-8) activity compared to females, whether this directly influenced the local cellular adaptations is unknown. Finally, the data suggest that leucine, HMB or BCAA supplementation did not influence muscle function following multiple eccentric contractions in females or males and these measures in females were not impacted by circulating oestradiol concentrations. Overall, there was no overarching difference between protein supplements following an acute bout of exercise, especially in reducing DOMS. Further research should focus including females as there were differences in CK and cytokine concentrations. Despite the occurrence of DOMS, the increased IL-7 with leucine, increased TNFr1-α with HMB and altered TNFr1-α with BCAA in females and males could provide improved adaptation in longer intervention studies especially in females and is an avenue for further investigation.

## Ethics Declarations

### Funding

Liverpool John Moores University.

### Conflicts of interest

No conflicts of interest.

### Ethical approval

The study involved human volunteers and was approved by Liverpool John Moores Research Ethics Committee (16/SPS/007), adhered to the Declaration of Helsinki and was registered respectively as a clinical trial (NCT04679519).

### Informed consent

All volunteers gave informed consent for this study.

### Author contributions

A.D.B designed the study, collected and analysed data, wrote and edited the manuscript. M.G, C.C.L.T, M.C.C collected and analysed data, commented on manuscript. C.E.S designed and supervised the study, provided funding and edited the manuscript.

## Notes

### Competing Interest Statement

The authors have declared no competing interest.

